# Sub-optimality of the early visual system explained through biologically plausible plasticity

**DOI:** 10.1101/799155

**Authors:** Tushar Chauhan, Timothée Masquelier, Benoit R. Cottereau

## Abstract

The early visual cortex is the site of crucial pre-processing for more complex, biologically relevant computations that drive perception and, ultimately, behaviour. This pre-processing is often viewed as an optimisation which enables the most efficient representation of visual input. However, measurements in monkey and cat suggest that receptive fields in the primary visual cortex are often noisy, blobby, and symmetrical, making them sub-optimal for operations such as edge-detection. We propose that this suboptimality occurs because the receptive fields do not emerge through a global minimisation of the generative error, but through locally operating biological mechanisms such as spike-timing dependent plasticity. Using an orientation discrimination paradigm, we show that while sub-optimal, such models offer a much better description of biology at multiple levels: single-cell, population coding, and perception. Taken together, our results underline the need to carefully consider the distinction between information-theoretic and biological notions of optimality in early sensorial populations.

## Introduction

The human visual system processes an enormous throughput of sensory data in successive operations to generate percepts and behaviours necessary for biological functioning (Anderson et al., 2005; Raichle, 2010). Computations in the early visual cortex are often explained through unsupervised normative models which, given an input dataset with statistics similar to our surroundings, carry out an optimisation of criteria such as energy consumption and information-theoretic efficiency (Bell & Sejnowski, 1997; Bruce et al., 2016; Hoyer & Hyvärinen, 2000; Olshausen & Field, 1996; van Hateren & van der Schaaf, 1998; Zhaoping, 2006). While such arguments do explain why many properties of the early visual system are closely related to characteristics of natural scenes (Bell & Sejnowski, 1997; Beyeler et al., 2019; Geisler, 2008; Hunter & Hibbard, 2015; Lee & Seung, 1999; Olshausen & Field, 1996), they are not equipped to answer questions such as how cortical structures which support complex computational operations implied by such optimisation may emerge, how these structures adapt, even in adulthood (Hübener & Bonhoeffer, 2014; Wandell & Smirnakis, 2010), and why some neurones possess receptive fields which are sub-optimal in terms of information processing (Jones & Palmer, 1987; Ringach, 2002).

It is now well established that locally-driven synaptic mechanisms such as spike-timing dependent plasticity (STDP) are natural processes which play a pivotal role in shaping the computational architecture of the brain (Brito & Gerstner, 2016; Caporale & Dan, 2008; Delorme et al., 2001; Larsen et al., 2010; Markram et al., 1997; Masquelier, 2012). Therefore, it is only natural to hypothesise that locally operating, biologically plausible models of plasticity must offer a better description of receptive fields in early visual cortex. However, such line of reasoning leads to the obvious question: what exactly constitutes a ‘better description’ of a biological system, and more specifically, the early visual cortex. Here, we use a series of criteria spanning across electrophysiology, information theory, and machine learning, to investigate how descriptions of early visual RFs provided by a local, abstract STDP model differ from two influential and widely employed normative schemes – Independent component analysis (ICA), and Sparse coding (SC). Our results demonstrate that a local process-based model of experience-driven plasticity is better suited to capturing the receptive fields (RFs) of simple-cells, thus suggesting that biological preference does not always concur with forms of global, information-theoretic optimality.

More specifically, we show that STDP units are able to better capture the characteristic sub-optimality in RF shape reported in literature (Jones & Palmer, 1987; Ringach, 2002), and their orientation tuning closely matches measurements in the macaque primary visual cortex (V1) (Ringach et al., 2002). To investigate the possible effects of this sub-optimality on downstream processing we estimated the performance of an ideal observer and a linear decoder on an orientation discrimination task. We find that decoding from the STDP population shows a bias for cardinal orientations (horizontal and vertical gratings) – something that is not observed for ICA and SC. This increase in discriminability for cardinal orientations has been reported in numerous behavioural studies (Heeley & Timney, 1988; Orban et al., 1984), and there is converging evidence which places its locus in the early visual cortex (Furmanski & Engel, 2000; Maffei & Campbell, 1970; Mansfield, 1974) – a claim that is amply supported by our results.

Taken together, our findings suggest that while the information carrying capacity of an STDP ensemble is not optimal when compared to generative schemes, it is precisely this sub-optimality which may make process-based, local models more suited for describing the initial stages of sensory processing.

## Results

We used an abstract model of the early visual system with three representative stages: retinal input, lateral geniculate nucleus (LGN) processing, and V1 activity (Figure 1B). To simulate retinal activity corresponding to natural inputs, patches of size 3° × 3° (visual angles) were sampled randomly from the Hunter-Hibbard database (Hunter & Hibbard, 2015) of natural scenes (Figure 1A). 10^5^ patches were used to train models corresponding to three encoding schemes: ICA, SC and STDP. Each model used a specific procedure for implementing the LGN processing and learning the synaptic weights between the LGN and V1 (see Figure 1B and *Methods*).

**Figure 1:**
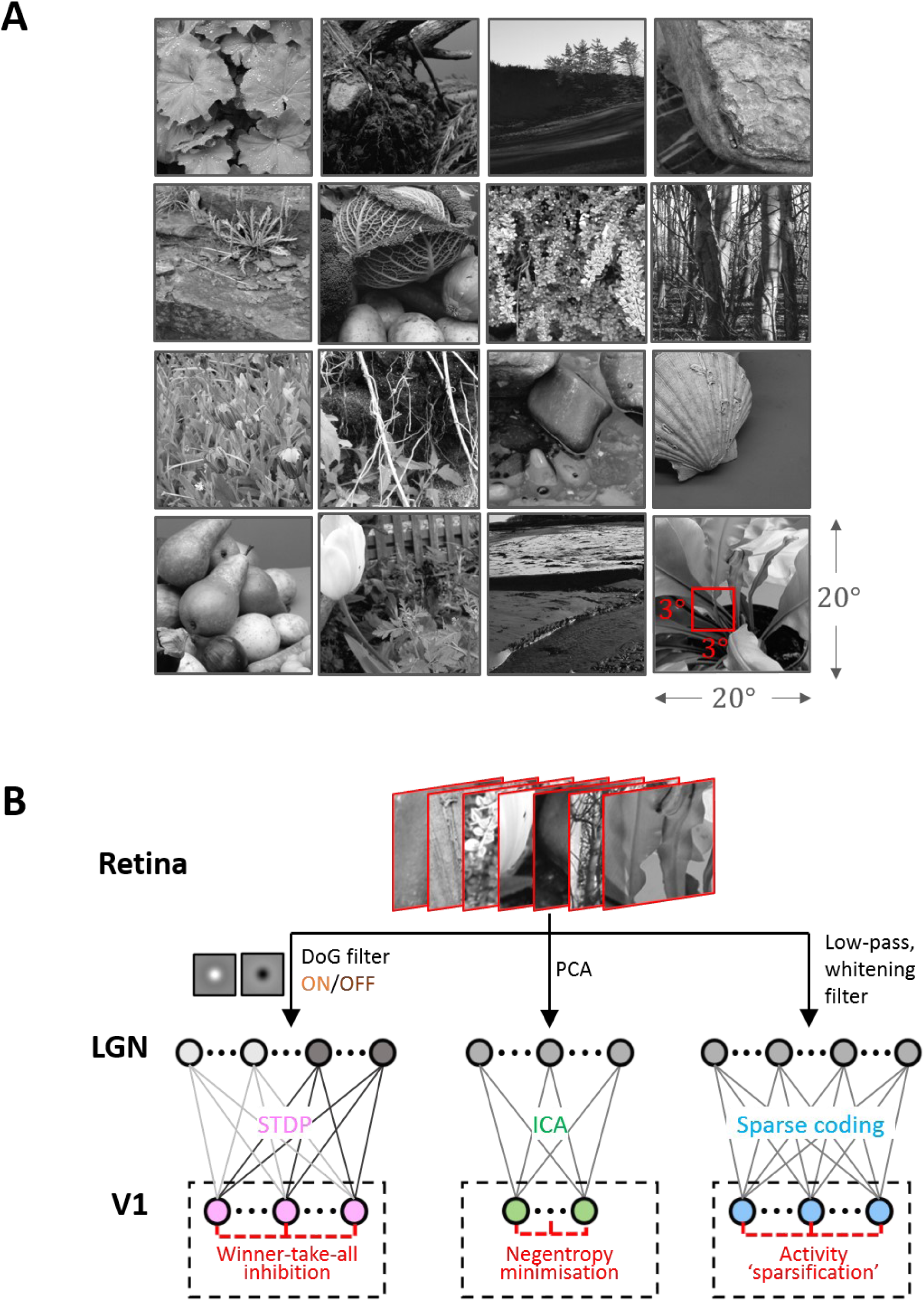
Dataset and the computational pipeline. **A.** Training data. The Hunter-Hibbard dataset of natural images was used. The images in the database have a 20° × 20° field of view. Patches of size 3° × 3° were sampled from random locations in the images (overlap allowed). The same set of 100,000 randomly sampled patches was used to train three models: Spike-timing dependent plasticity (STDP), Independent component analysis (ICA) and Sparse coding (SC). **B.** Modelling the early visual pathway. Three representative stages of early visual computation were captured by the models: retinal input, processing in the lateral geniculate nucleus (LGN), and the activity of early cortical populations in the primary visual cortex (V1). Each input patch represented a retinal input. This was followed by filtering operations generally associated with the LGN, such as decorrelation and whitening. Finally, the output from the LGN units/filters was connected to the V1 population through all-to-all (dense) plastic synapses which changed their weights during learning. Each model had a specific optimisation strategy for learning: the STDP model relied on a local rank-based Hebbian rule, ICA minimised mutual information (approximated by the negentropy), and SC enforced sparsity constraints on V1 activity. Abbreviations: DoG: Difference of Gaussian, PCA: Principal component analysis.

As expected, units in all models converged to oriented, edge-detector like RFs. While the RFs from ICA (Figure 2B) and SC (Figure 2C) were elongated and highly directional, STDP (Figure 2A) RFs were more compact and less sharply tuned. This is in agreement with simple-cell recordings in the macaque and cat, where studies have reported that not all RFs are optimally tuned for edge-detection (Jones & Palmer, 1987; Ringach, 2002). A quantitative measure of this phenomenon was obtained by fitting Gabor functions to the RFs and considering the frequency-normalised spread vectors or FSVs (Ringach, 2002) of the fit. The first component (*n*_*x*_) of the FSV characterises the number of lobes in the RF, and the second component (*n*_*y*_) is a measure of the elongation of the RF (perpendicular to carrier propagation). A considerable number of simple-cell RFs measured in macaque and cat tend to fall within the square bounded by *n*_*x*_ = 0.5 and *n*_*y*_ = 0.5. The FSVs of a sample of neurones (*N* = 93) measured in the macaque V1 (Ringach, 2002) indicate that 59.1% of the neurones lay within this region (Figure 2D). Since they are not elongated, and contain few lobes (typically 2-3 on/off regions), they tend to be compact – making them less effective as edge-detectors compared to more crisply tuned, elongated RFs. While a considerable number of STDP units (82.2%) tend to fall in this realistic regime, ICA (10.7%) and SC (4.0%) show a distinctive shift upwards and to the right.

**Figure 2:**
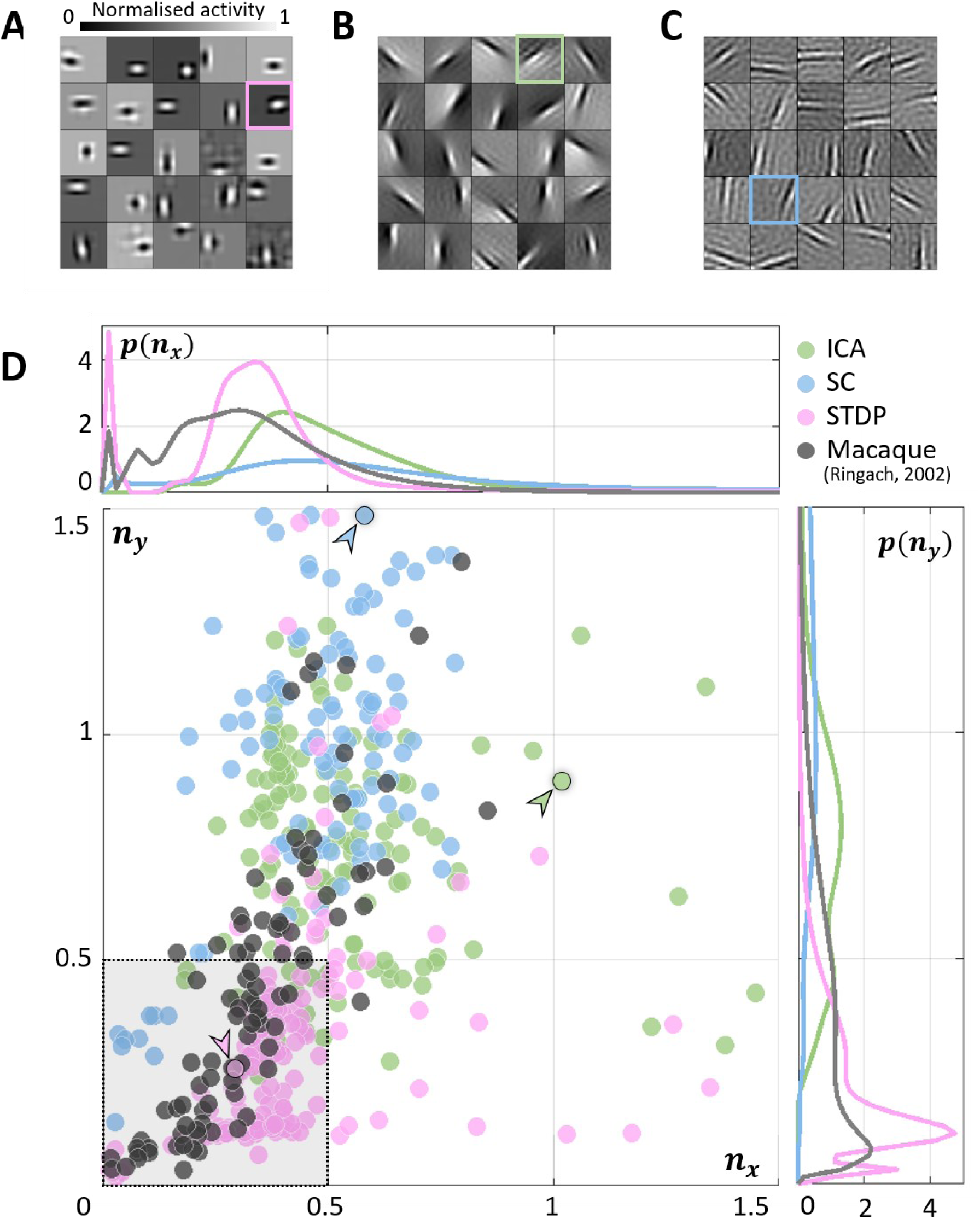
Receptive field (RF) shape. **A, B, C.** RFs of neurones randomly chosen from the three converged populations. The STDP population is shown in **A**, ICA in **B**, and SC in **C. D.** Frequency-scaled spread vectors (FSVs). FSV is a compact metric for quantifying RF shape. *n*_*x*_ is proportional to the number of lobes in the RF, *n*_*y*_ is a measure of the elongation of the RF, and values near zero characterise symmetric, often blobby RFs. The FSVs for STDP (pink), ICA (green) and SC (blue), are shown with data from macaque V1 (black) (Ringach, 2002). Measurements in macaque and cat simple-cells tend to fall within the square bound by 0.5 along both axes (shaded in grey, with a dotted outline). Three representative neurones are indicated by colour-coded arrows: one for each algorithm. The corresponding RFs are outlined in **A, B** and **C** using the corresponding colour. The STDP neurone has been chosen to illustrate a blobby RF, the ICA neurone shows a multi-lobed RF, and the SC neurone illustrates an elongated RF. Insets above and below the scatter plot show estimations of the probability density function for *n*_*x*_ and *n*_*y*_. Both axes have been cut-off at 1.5 to make comparison with biological data easier (complete distributions are shown in Supplementary Figure S 1).

The inlays in Figure 2D provide estimations of the probability densities of two FSV parameters for the macaque data and the three models. An interesting insight into these distributions is given by the Kullback-Leibler (KL) divergence from the model distributions to the distribution of the macaque data. KL divergence is a directed measure which can intuitively be interpreted as the additional number of bits required if one of the three models were used to encode data sampled from the macaque distribution. The KL divergence for the STDP model was found to be 3.0 bits indicating that, on average, it would require three extra bits to encode data sampled from the macaque distribution. In comparison, SC and ICA were found to require 8.4 and 14.6 additional bits respectively. An examination of the KL divergence of marginal distributions of the FSV parameters (Supplementary Table S 1) suggests that STDP offers a better encoding of both the *n*_*x*_ (number of lobes) and the *n*_*y*_ (compactness) parameters.

Given this sub-optimal, compact nature of STDP RF shapes, we next investigated how this affected the responses of these neurones to sharp edges. In particular, we were interested in how the orientation tuning curves of the units from the three models would compare to biological data. We simulated a typical electrophysiological experiment to probe orientation tuning (Figure 3A). To each unit, we presented noisy sine-wave gratings (SWGs) at its preferred frequency and recorded its activity as a function of the stimulus orientation. This allowed us to plot its orientation tuning curve (OTCs) (Figure 3B) and estimate the peak and tuning bandwidth. While the peak indicates the preferred orientation of the unit, the bandwidth is a measure of the local selectivity of the unit around its peak – low values corresponding to sharply tuned neurones and higher values corresponding to broadly tuned, less selective neurones. For all three models, we estimated the probability density functions of the OTC bandwidth, and compared it to the distribution estimated over a large set of data (*N* = 308) measured in macaque V1 (Ringach et al., 2002) (Figure 3C). The ICA and SC distributions peak at about 10-degrees (ICA: 9.1deg, SC: 8.5deg) while the STDP and macaque data have much broader tunings (STDP: 15.1deg, Macaque data: 19.1deg). This is also reflected in the KL divergence measured from the three model distributions to the macaque distribution (ICA: 2.4 bits, SC: 3.5 bits, STDP: 0.29 bits).

**Figure 3:**
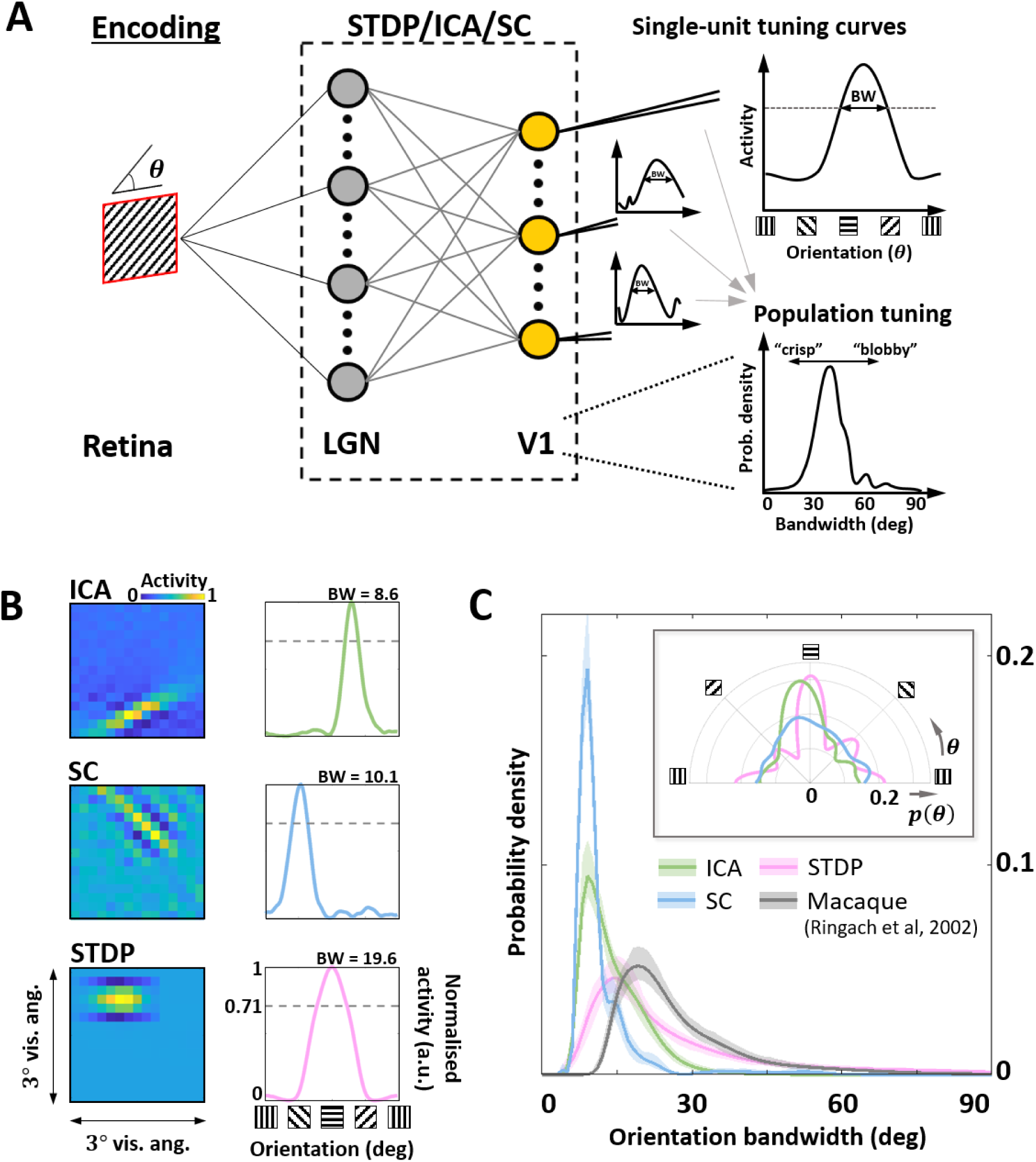
Orientation encoding. **A.** Orientation tuning. Sine-wave gratings with additive Gaussian noise (0dB SNR) were presented to the three models to obtain single-unit orientation tuning curves (OTCs). OTC peak identifies the preferred orientation of the unit, and OTC bandwidth (half width at 1/√2 peak response) is a measure of its selectivity around the peak. Low bandwidth values are indicative of sharply tuned units while high values signal broader, less specific tuning. **B.** Single-unit tuning curves. RF (left) and the corresponding OTC (right) for representative units from ICA (top row, green), SC (second row, blue) and STDP (bottom row, pink). The bandwidth is shown above the OTC. **C.** Population tuning. Estimated probability density of the OTC bandwidth for the three models (same colour code as panel **B**), and data measured in macaque V1 (black) (Ringach et al., 2002). The inlay shows estimated probability density of the preferred orientation for the three models.

After characterising the encoding capacity of the models, we next probed the possible downstream implications of such codes. The biological goal of most neural code, in the end, is the generation of behaviour that maximises evolutionary fitness. However, due to the complicated neural apparatus that separates behaviour from early sensory processing, it is not straightforward (or at times, even possible) to analyse the interaction between the two. Here, we make use of two separate metrics to investigate this relationship. In both cases, the models were presented with oriented SWGs, followed by an analysis of the resulting population activity (Figure 4A).

**Figure 4:**
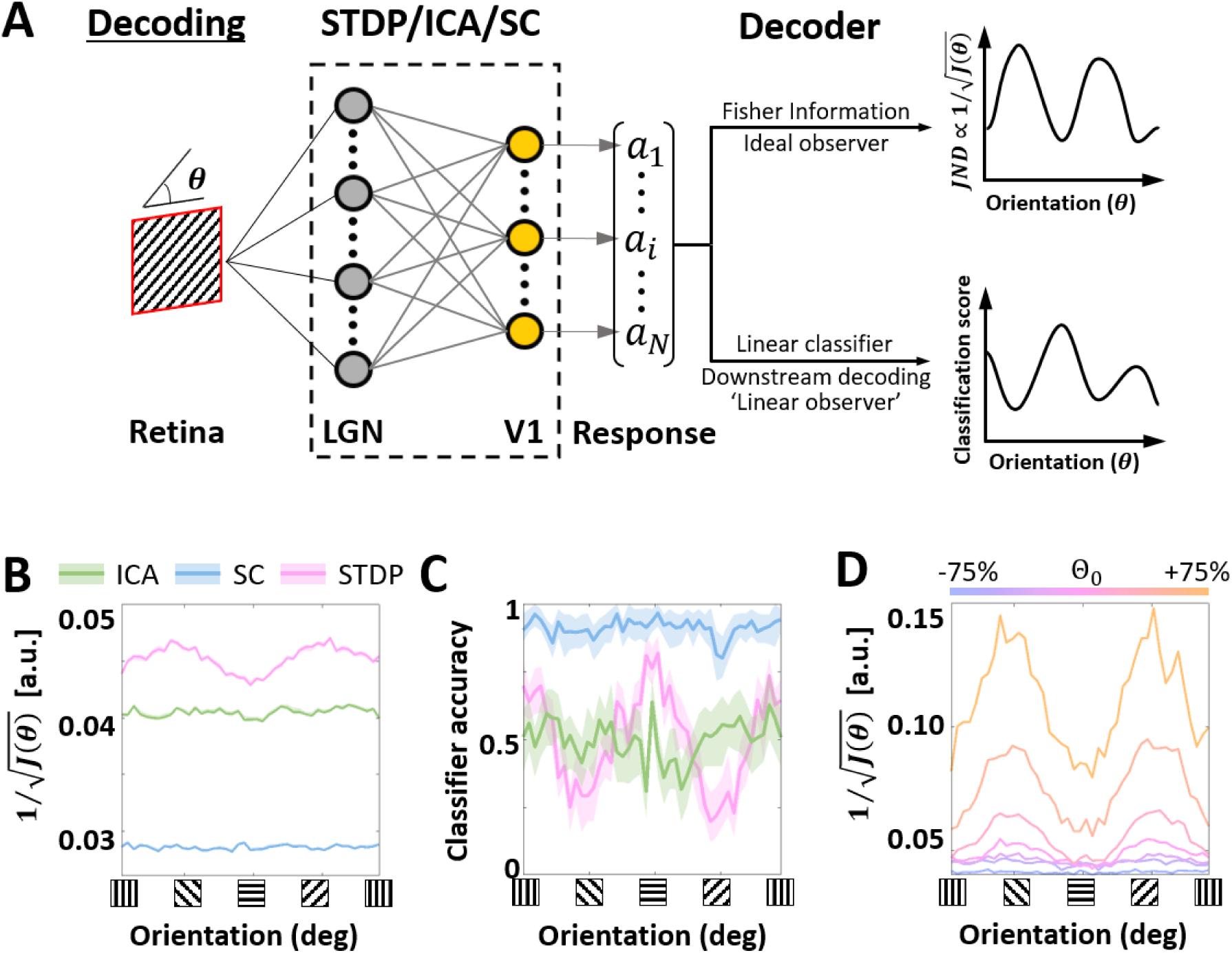
Orientation decoding. **A.** Retrieving encoded information. Sine-wave gratings (SWGs) with additive Gaussian noise at 0dB were presented to the three models. The following question was then posed: how much information about the stimulus (in this case, the orientation) can be decoded from the population responses? The theoretical limit of the accuracy of such a decoder can be approximated by estimating the Fisher information (FI) in the responses. This can be interpreted as the response of an ideal observer capable of extracting all the information contained in the population responses. In addition, a linear decoder was also used to directly decode the population responses. This could be a downstream process which is linearly driven by the population activity, or a less-than-optimal ‘linear observer’. **B.** Ideal observer (Fisher information). The abscissa shows the stimulus orientation, and the ordinate shows the inverse square-root of the estimated FI. Due to the Cramér-Rao bound, low values on the ordinate denote more precise encoding of the orientation, while higher values denote lower precision. Results for ICA are shown in green, SC in blue, and STDP in pink. **C.** Linear decoding. The stimuli and responses from **B** were used to train a linear-discriminant classifier. The ordinate shows the accuracy (probability of correct classification) for each ground-truth value of the stimulus orientation (abscissa). The same symbols as **B** are used, and envelopes around the solid lines show 95% confidence intervals. **D.** Post-training threshold variation in STDP. The SWG stimuli used in **C** were used to test STDP models with different values of the threshold parameter, which represents the threshold potential of the neurones in the model. The threshold was either increased (by 25%, 50% or 75%) or decreased (by 25%, 50% or 75%) with respect to the training threshold (denoted by Θ_0_). The abscissa and ordinate denote the same quantities as **B**, and each line shows the ideal observer response from one of the models. The magnitude of change in the threshold is denoted by a corresponding change in colour from pink (Θ_0_) to orange (increase) or blue (decrease).

In the first analysis, we estimated the Fisher information (FI) contained in the generated responses. Orientation discrimination tasks are particularly suited to such an analysis (i.e., the study of a behavioural measure using early sensory activity) as V1 activity has been shown to correlate with orientation discrimination thresholds (Vogels & Orban, 1990), something that has not been observed for other visual properties such as binocular disparity (Nienborg & Cumming, 2006) or motion (Grunewald et al., 2002). Given the responses of an encoding model, the Crámer-Rao bound permits us to use the FI to draw inferences about the optimal discrimination performance – leading to the simulation of what could be called a locally-optimal ideal observer (Wei & Stocker, 2017). We find that ICA and SC ideal observers have lower thresholds (i.e., a better orientation discrimination performance) compared to STDP (Figure 4B). Considering the sharper population tuning of the ICA and SC models, this is not surprising. Interestingly, the STDP ideal observer shows a bias for cardinal orientations (horizontal and vertical gratings), while ICA and SC ideal observers show uniform performance across all stimulus orientations.

Although the FI ideal observer represents the optimal decoding of a given encoding scheme, it does not automatically follow that all the encoded information is available for downstream processing (Quiroga & Panzeri, 2009). This is especially true in the presence of significant higher-order correlations (Shamir & Sompolinsky, 2004). To investigate more realistic orientation decoding in the three models, we implemented a second analysis where we reduced the complexity of the decoding and examined the performance of a decoder built on linear discriminant classifiers (these classifiers assume fixed first-order correlations in the input). Such decoders could be interpreted as linearly-driven feedforward populations downstream from the thalamo-recipient layer (the ‘V1’ populations in the three models), or a simplified, ‘linear’ observer. In general, we found the decoding performance (Figure 4C) to be consistent with ideal observer predictions (Figure 4B), indicating that linear correlations represent a sizeable amount of correlations in all three encoding schemes. Both ICA and SC showed uniform decoding scores across the orientations, with SC being the most accurate of the three models. Once again, STDP showed a modulation at the cardinal orientations, with the scores being almost three times higher at the peaks (horizontal and vertical gratings) than the troughs.

Since the STDP model is linear up to a thresholding operation, we hypothesised that the modulation in decoding (Figure 4A and B) must result from the intervening thresholding nonlinearity. To test this hypothesis, we ran simulations where the threshold parameter (analogue of the threshold potential in the STDP model) was either increased or decreased. Note that in all simulations the model was first trained (i.e., synaptic learning using natural stimuli, see Figure 1) using the same ‘training’ threshold, and the increase/decrease of the threshold parameter was imposed post-convergence. Our results (Figure 4D) showed that the modulation of the classification performance for horizontal and vertical gratings became more pronounced as the threshold was increased, and flattened as the threshold was decreased. However, this increase in the cardinal orientation bias came at the cost of an overall decrease in precision.

## Discussion

Traditionally, process-based descriptions (often studied through mean-field statistics under large *N* limits) have been used to model fine-grained neuronal dynamics (Harnack et al., 2015; Kang & Sompolinsky, 2001; Moreno-Bote et al., 2014), while more global, normative schemes are employed to predict population-level characteristics (Hoyer & Hyvärinen, 2000; Lee & Seung, 1999; Olshausen & Field, 1997; van Hateren & van der Schaaf, 1998). Detailed process-based models suffer from constraints imposed by computational complexity, prohibitively long execution times (which do not scale well for large networks), and hardware that is geared towards synchronous processing. On the other hand, most one-shot models can leverage faster computational libraries and architectures developed over decades, thereby leading to more efficient and scalable computation. Through this work, we argue that more abstract forms of process-based models, when used to answer specific questions, can still give a closer approximation of biological processes than normative schemes. Our model, despite important limitations such as the use of at most one spike per neuron, lack of cortico-cortical connections, and feedback from higher visual areas, is still able to describe a number of phenomena which are not predicted by normative schemes.

The converged RFs of the STDP model are closer to those reported in electrophysiological measurements in the macaque. A number of them display a characteristic departure from the optimal, sharply tuned edge-detectors predicted by ICA and SC (Figure 2). This suboptimal shape also results in suboptimal orientation encoding, which, once again, was found to be closer to data measured in the macaque V1 (Figure 3). Furthermore, decoding from this population shows a distinct bias for horizontal and vertical stimuli, with the performance for the cardinal orientations being roughly two times better than other orientations (Figure 4). Interestingly, this bias for cardinal directions has also been observed in human participants (Orban et al., 1984), who also show an almost two-fold reduction in just-noticeable-differences for horizontal and vertical stimuli (Heeley & Timney, 1988). Our results suggest that such behavioural biases for cardinal directions (the so-called oblique effect) are supported by the cortical architecture as early as the early thalamocortical interface – a claim that is further reinforced by electrophysiology (Dragoi et al., 2001; Li et al., 2003) and imaging experiments (Furmanski & Engel, 2000) which suggest that neural activity in the primary visual cortex may be directly correlated with the perceptual oblique effect.

Moreover, our simulations also demonstrated that the magnitude of the cardinal orientation bias in the STDP model can be changed by manipulating the spiking threshold (Figure 4D). Thus, even after the initial learning phase has ended, the network is still theoretically capable of shrinking or expanding its encoded dictionary without modifying its synaptic connectivity – simply by modulating its threshold. In the cortex, such changes are likely to be mediated by homeostatic processes (Harnack et al., 2015; Perrinet, 2019) and metaplastic regulation (Abraham, 2008; Cooke & Bear, 2010). This supports the idea that purely bottom-up changes are indeed capable of driving perceptually relevant cortical changes in adults who are beyond the critical window of plasticity – an observation of particularly relevance for reports claiming that low-level changes in the early visual cortex may accompany perceptual learning (Bao et al., 2010; Furmanski et al., 2004). In the context of the current model, this is only possible if the information content in the non-spiking activity is larger than the spiking activity (Figure S 5), and suggests that the model may employ some form of variable-length encoding. This becomes especially pertinent for realistic natural inputs which are more complicated and varied than SWGs. When we characterised the sparsity of non-spiking activity for each of the three models using naturalistic stimuli (Figure 5A), our results (Figure 5B) confirmed that the STDP responses do indeed show a higher variability in sparsity when measured across both the encoding ensemble (‘spatial sparsity’), and the input sequence (‘temporal sparsity’). Thus, the sparsity of the STDP neurones is stimulus-dependent, and likely driven by the relative probability of occurrence of specific features in the dataset.

**Figure 5:**
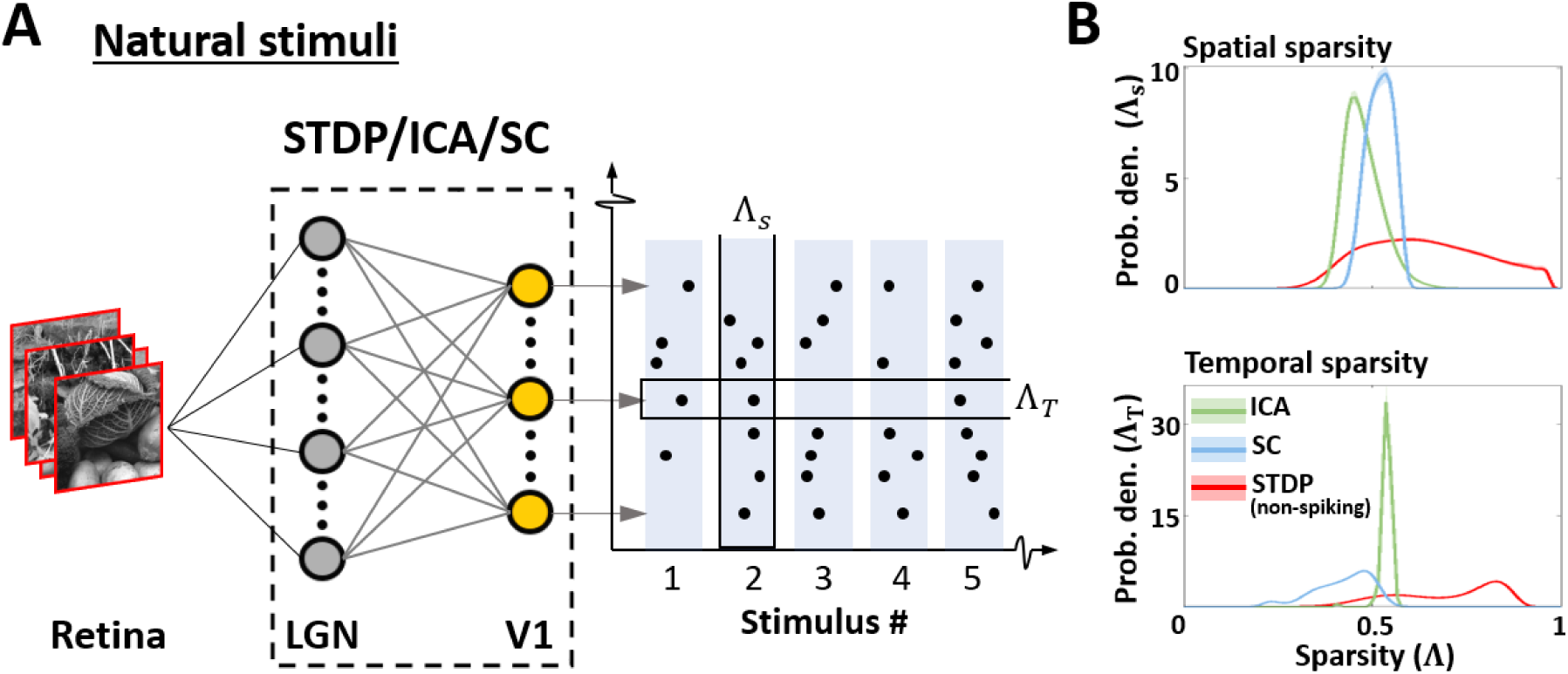
Sparsity of responses to natural stimuli. **A.** Sparsity indices. To estimate the sparsity of the non-spiking responses to natural stimuli, 10^4^ patches (3° × 3° visual angle) randomly sampled from natural scenes were presented to the three models. Two measures of sparsity were defined: Spatial sparsity Index (Λ_*S*_) was defined as the average sparsity of the activity of the entire neuronal ensemble, while Temporal sparsity Index (Λ_*T*_) was defined as the average sparsity of the activity of single neurones to the entire input sequence. **B.** Spatial and temporal sparsity. The top panel shows the estimated probability density of Λ_*S*_ for ICA (green), Sparse coding (blue) and STDP (red). Λ_*S*_ varied between 0 (all units activate with equal intensity) and 1 (only one unit activates) by definition. The bottom panel shows the estimated probability density of Λ_*T*_ in a manner analogous to Λ_*S*_. Λ_*T*_ also varied between 0 (homogeneous activity for the entire input sequence) and 1 (activity only for few inputs in the sequence).

The encoding that emerges from localised learning, as we have shown, can confer biological advantages while being sub-optimal in a global information-theoretic framework. This dichotomy has indeed been characterised at various stages of the early visual system (Chelaru & Dragoi, 2008; Field & Chichilnisky, 2007; Liu et al., 2009; Ringach, 2002; Ringach et al., 2002), and offers an interesting glimpse into the notion of ‘optimality’ in biological systems. In such systems input information is not the only criterion of fitness, and the landscape is modulated by factors such as the topological and dynamical constraints of the learning mechanisms, decoding capabilities of the downstream apparatus, and relevance of the resulting behaviour(s) to the functioning and survival of the organism. Given these realistic and complicated constraints, it is obvious that a clear global optimisation function which describes the responses of real neural populations is very difficult, if not impossible, to formulate. This makes an ideal case for the use of process-based, local descriptions which, by definition, offer a much deeper insight into the organisation and functioning of biological systems. Such models offer the flexibility to not only describe normative hypotheses about stimulus encoding, but to also predict how meaningful internal parameters and interactions in the model impact the behaviour of the system. With increasingly common availability of faster and more adaptable computing solutions, we hope process-based modelling will be adopted more widely by cognitive and computational neuroscientists.

Finally, it must be noted that in this study we have used classical forms of the ICA and SC algorithms. More generally, these models are part of a class of data-adaptive encoding schemes which also comprises other algorithms such as principal-component analysis (PCA) and nonnegative matrix factorisation (NMF). These algorithms rely on the optimality of the encoding manifold at representing the training data, and their use in neuroscience is motivated by the proven energetic and information-theoretic efficiency of neural processes at representing natural statistics. We would like to draw attention to a growing number of insightful studies based on hybrid encoding schemes which address multiple optimisation criteria (Beyeler et al., 2019; Martinez-Garcia et al., 2017; Perrinet & Bednar, 2015), often through local process-based computation (Isomura & Toyoizumi, 2018; Savin et al., 2010; Zylberberg et al., 2011).

## Methods

### Dataset

The Hunter-Hibbard dataset of natural images was used (Hunter & Hibbard, 2015). It is available under the MIT license at https://github.com/DavidWilliamHunter/Bivis, and consists of 139 stereoscopic images of natural scenes captured using a realistic acquisition geometry and a 20 degree field of view. Only images from the left channel were used, and each image was resized to a resolution of 5 pixels/degree along both horizontal and vertical directions. Inputs to all encoding schemes were 3 × 3 degree patches (i.e. 15 × 15 pixels) sampled randomly from the dataset (Figure 1A).

### Encoding models

Samples from the dataset were used to train and test three models corresponding to ICA, SC and STDP encoding schemes. Each model consisted of three successive stages (Figure 1B). The first stage represented retinal activations. This was followed by a pre-processing stage implementing operations which are typically associated with processing in the LGN, such as whitening and decorrelation. In the third stage, LGN output was used to drive a representative V1 layer.

During learning, 10^5^ patches (3 × 3 degree) were randomly sampled from the dataset to simulate input from naturalistic scenes. In this phase, the connections between the LGN and V1 layers were plastic, and modified in accordance with one of the three encoding schemes. Care was taken to ensure that the sequence of inputs during learning was the same for all three models. After training, the weights between the LGN and V1 layers were no longer allowed to change. The implementation details of the three models are described below.

#### Sparse Coding

SC algorithms are based on energy-minimisation, which is typically achieved by a ‘sparsification’ of activity in the encoding population. We used a now-classical SC scheme proposed by (Olshausen & Field, 1996, 1997). The pre-processing in this algorithm consists of an initial whitening of the input using low pass filtering, followed by a trimming of higher frequencies. The latter was employed to counter artefacts introduced by high frequency noise, and the effects of sampling across a uniform square grid. In the frequency domain the pre-processing filter was given by a zero-phase kernel:

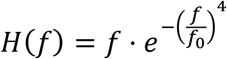

Here, *f*_0_ = 10 cycles/degree is the cut-off frequency. The outputs of these LGN filters were then used as inputs to the V1 layer composed of 225 units (3° × 3° RF at 5 pixels/degree). Retinal projections of the converged RFs (Figure 2C) were recovered by an approximate reverse-correlation algorithm (Ringach, 2002; Ringach & Shapley, 2004) derived from a linear-stability analysis of the SC objective function about its operating point. The RFs (denoted as columns of a matrix, say ***ξ***) were given by:

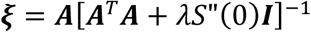

Here, ***A*** is the matrix containing converged sparse components as column vectors, *λ* is the regularisation parameter (for the reconstruction, it is set to 0.14*σ*, where *σ*^2^ is the variance in the input dataset), and *S*(*x*) is the shape-function for the prior distribution of the sparse coefficients (this implementation uses *log* (1 + *x*^2^)).

#### Independent Component Analysis (ICA)

ICA algorithms are based on the idea that the activity of an encoding ensemble must be as information-rich as possible. This typically involves a maximisation of mutual information between the retinal input and the activity of the encoding ensemble. We used a classical ICA algorithm called *fastICA* (Hyvärinen & Oja, 2000) which achieves this through an iterative estimation of input negentropy. The pre-processing in this implementation was performed using a truncated Principal Component Analysis (PCA) transform (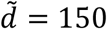 components were used), leading to low-pass filtering and local decorrelation akin to centre-surround processing reported in the LGN. If the input patches are denoted by the columns of a matrix (say ***X***), the LGN activity ***L*** can be written as:

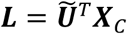

Here, ***X***_*C*_ = ***X*** − ⟨***X***⟩ and ***Ũ*** is a matrix composed of the first 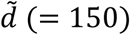 principal components of ***X***_*C*_. The activity of these LGN filters was then used to drive the ICA V1 layer consisting of 150 units, with its activity **Σ** being given by:

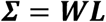

Here, ***W*** is the un-mixing matrix which is optimised during learning. The recovery of the RFs for ICA was relatively straight forward, as, in our implementation, they were assumed to be equivalent to the filters which must be applied to a given input to generate the corresponding V1 activity. The RFs (denoted as columns of a matrix, say ***ξ***) were given by:

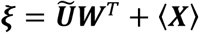

#### Spike timing dependent plasticity (STDP)

STDP is a biologically observed, Hebbian-like learning rule which relies on local spatiotemporal patterns in the input. We used a feedforward model based on an abstract rank-based STDP rule (Chauhan et al., 2018). The pre-processing in the model consisted of ON/OFF filtering using difference-of-Gaussian filters based on the properties of magno-cellular LGN cells. The outputs of these filters were converted to relative first-spike latencies using a monotonically decreasing function (1/*x* was used), and only the earliest 10% spikes were allowed to propagate to V1 (Delorme et al., 2001; Masquelier & Thorpe, 2007). These spikes were used to drive an unsupervised network of 225 (non-leaky) integrate-and-fire neurones. During learning, changes in the synaptic weights between LGN and V1 were governed by a simplified version of the STDP rule proposed by (Gütig et al., 2003). After each iteration, the change (Δ*w*) in the weight (*w*) of a given synapse was given by:

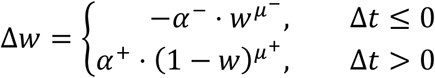

Here, *Δt* is the difference between the post- and pre-synaptic spike times, and the constants *α*^±^ and *µ*^±^ describe the learning-rate and non-linearity of the process respectively. Thus, the weight was increased if a presynaptic spike occurred before the postsynaptic spike (causal firing), and decreased if it occurred after the post-synaptic spike (acausal firing). During each iteration of learning, the population followed a winner-take-all inhibition rule wherein the firing of one neurone reset the membrane potentials of all other neurones. After learning, this inhibition was no longer active and multiple units were allowed to fire for each input. The RFs of the converged neurones were recovered using a linear approximation. If *w*_*i*_ denotes the weight of the synapse connecting a given neurone to the *i*th LGN filter, the RF ***ξ*** of the neurone was given by:

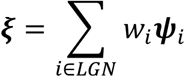

Here, ***ψ***_*i*_ is the RF of the *i*th LGN neurone.

### Evaluation metrics

#### Gabor fitting

Linear approximations of RFs (Figure 2, panels A, B and C) obtained by each encoding strategy were fitted using 2-D Gabor functions. This is motivated by the fact that all the encoding schemes considered here lead to linear, simple-cell-like RFs. In this case, the goodness-of-fit parameter (*R*^2^) provides an intuitive measure of how Gabor-like a given RF is. The fitting was carried out using an adapted version of the code available here, generously shared under an open-source license by Gerrit Ecke (University of Tübingen).

#### Frequency-normalised spread vector

The shape of the RFs approximated by each encoding strategy was characterised using frequency-normalised spread vectors (FSVs) (Chauhan et al., 2018; Ringach, 2002). For a RF fitted by a Gabor-function with sinusoid carrier frequency *f* and envelope size *σ* = [*σ*_*x*_ *σ*_*y*_]^T^, the FSV is given by:

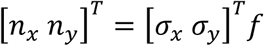

While *n*_*x*_ provides an intuition of the number of cycles in the RF, *n*_*y*_ is a cycle-adjusted measure of the elongation of the RF perpendicular to the direction of sinusoid propagation. The FSV serves as a compact, intuitive descriptor of the RF shape-invariance to affine operations such as translation, rotation and isotropic scaling.

#### Fisher Information

The information content in the activity of the converged units was quantified by using approximations of the Fisher information (FI, denoted here by the symbol Φ). If ***x*** = {*x*_1_, *x*_2_, *x*_3_, …, *x*_*N*_} is a random variable describing the activity of an ensemble of *N* independent units, the FI of the population with respect to a parameter *θ* is given by:

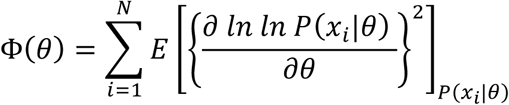

Here, 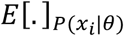 denotes expectation value with respect to the firing-state probabilities of the *i*^*th*^ neurone in response to the stimuli corresponding to parameter value *θ*. In our simulations, *θ* was the orientation (defined as the direction of travel) of a set of sine-wave gratings (SWGs) with additive Gaussian noise. The SWGs were presented at frequency of 1.25cycles/visual degree, and the responses were calculated by averaging over 8 evenly spaced phase values in [0°, 360°). This effectively simulated a drifting grating design within the constraints of the computational models. In Figure 4 (panels B and D) we report values for an SNR of 0dB (i.e., the signal and noise had equal variance – a fairly noisy condition), but the behaviour of the system was found to be similar for SNR values of −3dB and 3dB (Supplementary Figure S 2). The ground-truth values of *θ* were sampled at intervals of 4° in the range [0°, 180°), and each simulation was repeated 100 times.

#### Decoding using a linear classifier

In addition to FI approximations, we also used a linear decoder on the population responses obtained in the FI analysis. The decoder was an error-correcting output codes model composed of binary linear-discriminant classifiers configured in a one-vs.-all scheme. Similar to the FI experiment, ground-truth values of the orientation at intervals of 4° in the range [0°, 180°) were used as the class labels, and the activity generated by the corresponding SWG stimuli with added Gaussian noise was used as the training/testing data. The SWGs were presented at frequency of 1.25cycles/visual degree, and the responses were calculated by averaging over 8 evenly spaced phase values in [0°, 360°). Each simulation was repeated 100 times, each time with 5-fold validation. Figure 4C shows the results for a 0dB SNR, but the overall trends remained unchanged for −3dB and 3dB SNR levels (Supplementary Figure S 3).

#### Post-convergence threshold variation in STDP

To test how post-learning changes in the threshold affect the specificity of a converged network, we tested an STDP network trained using a threshold *Θ*_*training*_ by increasing or decreasing its threshold (to say, *Θ*_*testing*_) and presenting it with SWGs (same stimuli as the ones used to calculate the FI). We report the results of seven simulations where the relative change in threshold was given by 25% increments/decrements, i.e.:

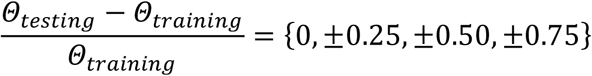

The results reported in Figure 4D were calculated for an input SNR of 0dB, but similar results were obtained for −3 and 3dB SNR as well (Supplementary Figure S 4).

#### Kullback-Leibler divergence

For each model, we estimated probability density functions (pdfs) over parameters such as the FSVs and the population bandwidth. To quantify how close the model pdfs were to those estimated from the macaque data, we employed the Kullback-Leibler (KL) divergence. KL divergence is a directional measure of distance between two probability distributions. Given two distributions *P* and *Q* with corresponding probability densities *p* and *q*, the KL divergence (denoted *D*_*KL*_) from a *Q* to *P* is given by:

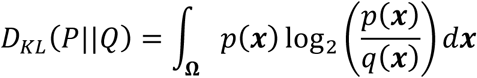

Here, **Ω** is the support of the distribution *Q*. In our analysis, we considered *p* as a pdf estimated from the macaque data, and *q* as the pdf (of the same variable) estimated using ICA, SC or STDP. In this case, KL divergence lends itself to a very intuitive interpretation: it can be considered as the additional bandwidth (in bits) which would be required if the biological variable were to be encoded using one of the three computational models. Note that *P* and *Q* may be multivariate distributions.

#### Sparsity: Gini index

The sparseness of the encoding was evaluated using the Gini index (GI). GI is a measure which characterises the deviation of the population-response from a uniform distribution of activity across the samples. It is 0 if all units have the same response and tends to 1 as the responses become sparser (being equal to 1 if only one unit responds, while others are silent). It is invariant to the range of the responses within a given sample, and robust to variations in sample-size (Hurley & Rickard, 2009). Formally, the GI (denoted here as Λ) is given by:

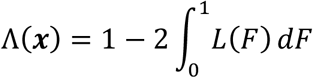

Here, *L* is the Lorenz function defined on the cumulative probability distribution *F* of the neural activity (say, ***x***). We defined two variants of the GI which measure the spatial (Λ_s_) and temporal sparsity (Λ_T_) of an ensemble of encoders (Figure 5A). Given a sequence of *M* inputs to an ensemble of *N* neurones, the spatial sparsity of the ensemble response to the *m*th stimulus is given by:

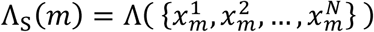

Here, 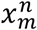 denotes the activity of the *n*th neurone in response to the *m*th input. Similarly, the temporal sparsity of the *n*th neurone over the entire sequence of inputs is given by:

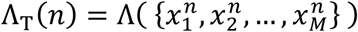

### Code

The code for ICA was written in python using the sklearn library which implements the classical *fastICA* algorithm. The code for SC was based on the C++ and Matlab code shared by Prof. Bruno Olshaussen. The STDP code was based on a previously published binocular-STDP algorithm available here.

## Acknowledgements

We are grateful to Dr. Dario Ringach for generously sharing single-cell data, and to Dr. Bruno Olshausen for sharing the original/initial versions of his sparse-coding routines. We would also like to thank Dr. Yves Trotter and Dr. Lionel Nowak for their critical feedback.

## Supplementary Material

### Kullback-Leibler divergence of receptive-field shape distributions

**Table S 1:**
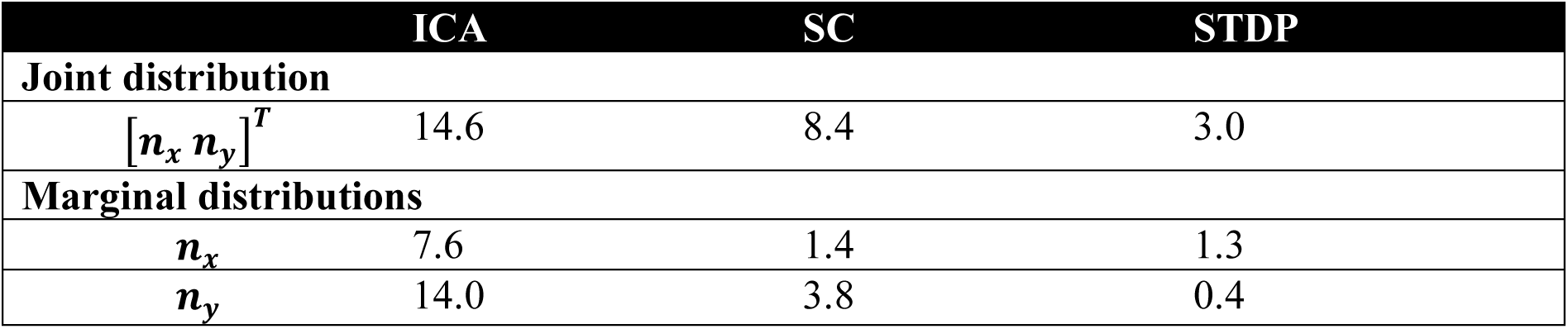
Kullback-Leibler (KL) divergence of frequency-normalised spread vectors (FSVs) from the models to the macaque distribution. The receptive-field (RF) shape of the neurones from the models and measurements in macaque V1 (Ringach, 2002) was parametrised by estimating the frequency-normalised spread vectors (FSVs). FSVs are characterised by two parameters *n*_*x*_ and *n*_*y*_: *n*_*x*_ is proportional to the number of lobes in the receptive field, and *n*_*y*_ is modulated by its elongation. The KL divergence reflects the number of additional bits required to encode the parameter(s) of interest from the macaque data using the distributions from one of the three models (ICA, SC or STDP). The KL divergence for the FSV reported in the Results section was based on an estimation of the joint distribution of the two FSV parameters. Here, we also report the KL divergence for the marginal distribution of each FSV parameter separately. All values are in bits.

|  | ICA | SC | STDP |
| --- | --- | --- | --- |
| **Joint distribution** |  |  |  |
| $[n_x \ n_y]^T$ | 14.6 | 8.4 | 3.0 |
| **Marginal distributions** |  |  |  |
| $n_x$ | 7.6 | 1.4 | 1.3 |
| $n_y$ | 14.0 | 3.8 | 0.4 |

### Frequency-normalised spread vector distributions

**Figure S 1:**
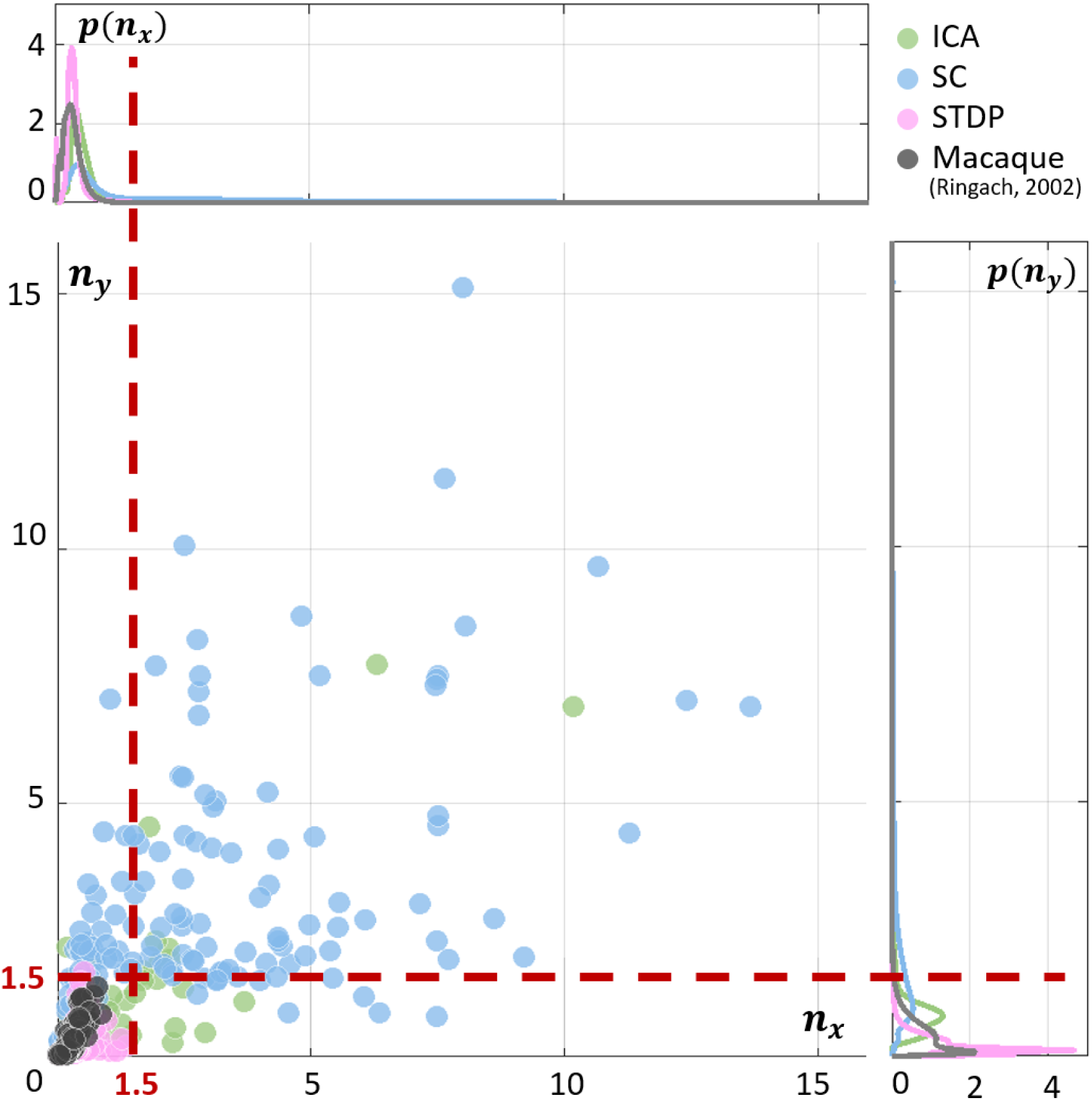
Frequency-normalised spread vector (FSV) distributions. FSVs are a compact metric for quantifying RF shape. To facilitate comparison with macaque data in Figure 2D, the two axes were cut off at 1.5 (marked with red, dashed lines). Here, we show the data without any cut-offs. The macaque data (black) and STDP (pink) are limited to a small region near the origin, while ICA (green) and SC (blue) show a much higher range of values.

### Effect of noise on ideal observer responses

**Figure S 2:**
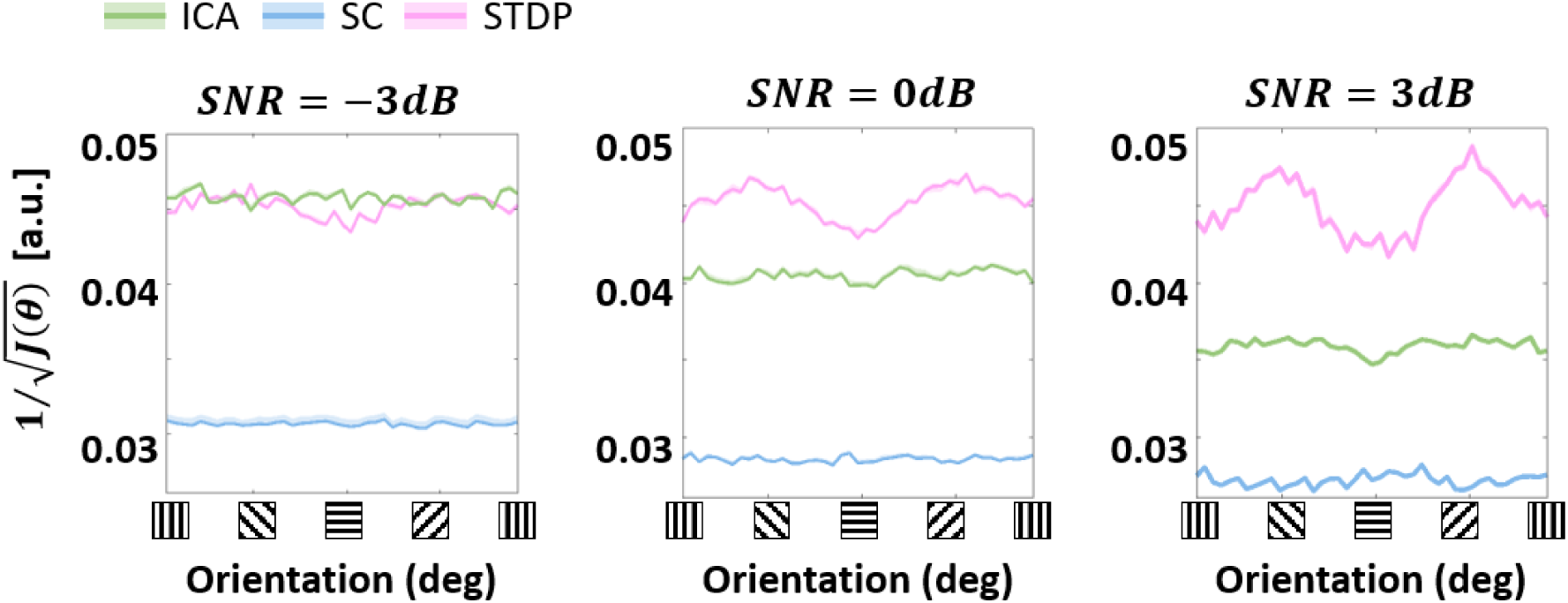
Ideal observer response across noise levels. Approximations of Fisher information were used to estimate ideal observer responses to sine-wave gratings with additive Gaussian noise (Figure 4B). The approximations were made at three noise levels: −3dB, 0dB and 3dB, corresponding to the variance of the noise being double, equal to or half the signal variance.

### Effect of noise on linear decoding

**Figure S 3:**
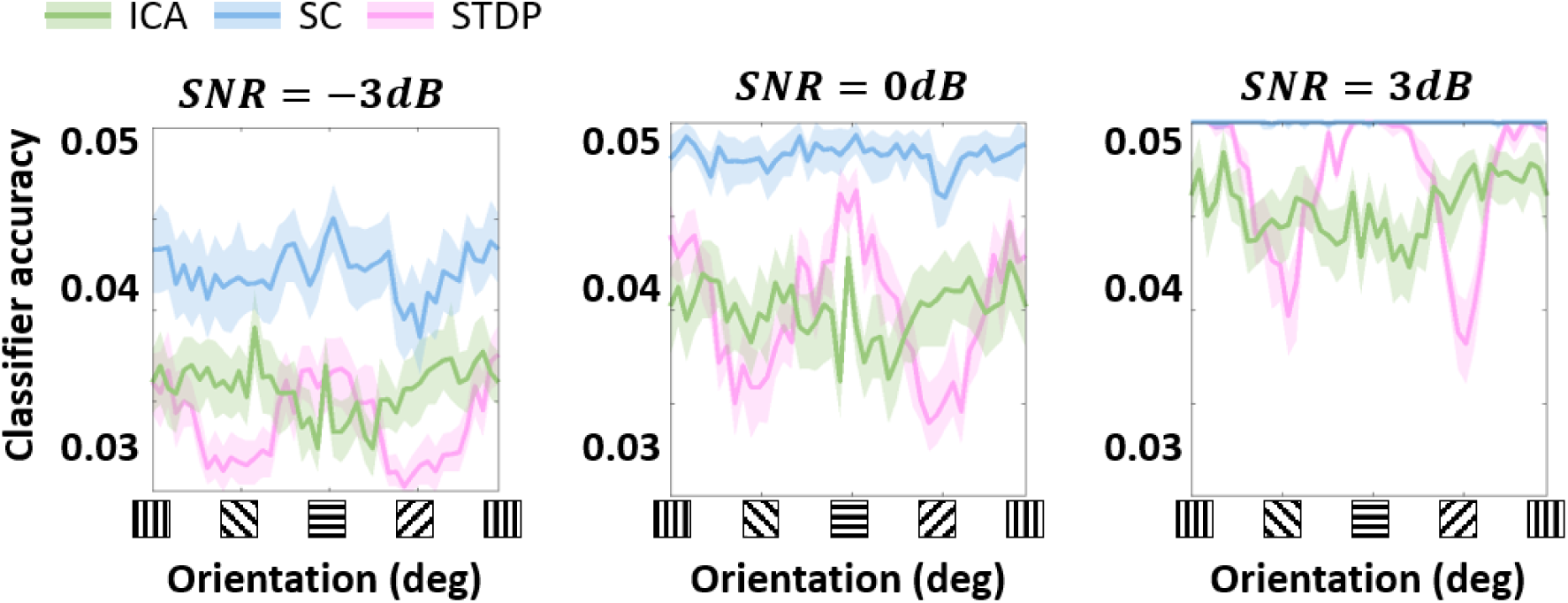
Classification accuracy across noise levels. Decoding scores of a linear classification model were calculated by using stimulus orientation as the class labels and population responses to the stimuli as the training/testing data (Figure 4C). The stimuli used were sine-wave gratings with three levels of additive Gaussian noise: −3dB, 0dB and 3dB, corresponding to the variance of the noise being double, equal to or half the signal variance.

### Effect of noise on the cardinal orientation bias

**Figure S 4:**
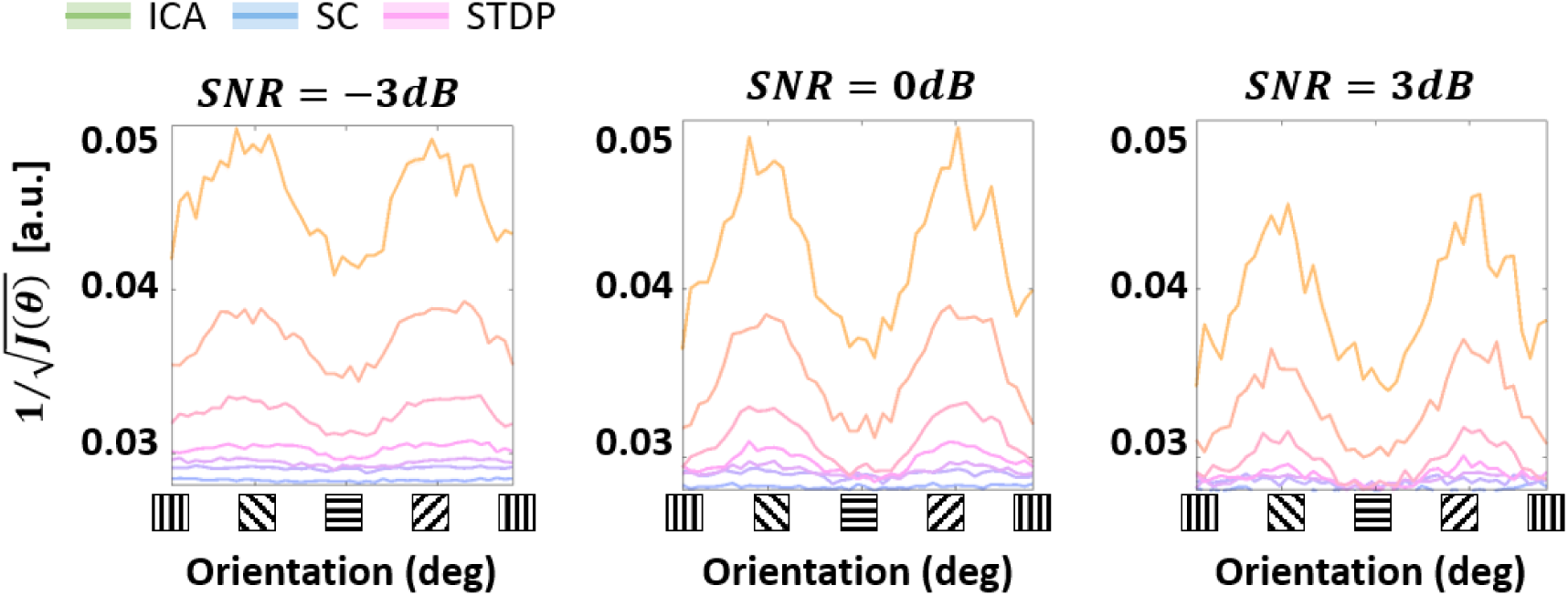
Cardinal orientation bias across noise levels. The threshold of the STDP model was increased/decreased post-convergence to investigate its effect on the modulation of the ideal observer response at horizontal and vertical (cardinal) orientations (Figure 4D). Oriented sine-wave gratings with additive Gaussian noise were used for these simulations. The simulations were run at three noise levels: −3dB, 0dB and 3dB, corresponding to the variance of the noise being double, equal to or half the signal variance.

### Fisher information from spiking and non-spiking responses

**Figure S 5:**
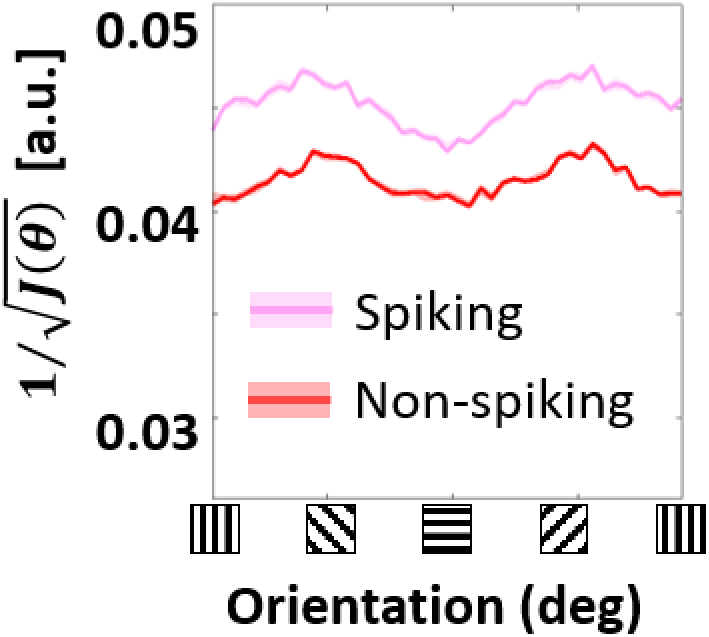
Spiking and non-spiking activity. The STDP model was presented with oriented sine-wave grating stimuli (see Methods and Figure 4A) and the resulting activity was used to estimate the Fisher information using spiking and non-spiking responses of the network. This was used to compute ideal-observer performance as a function of the stimulus orientation.

